# The mechanical stability of the world’s tallest broadleaf trees

**DOI:** 10.1101/664292

**Authors:** T. Jackson, A. Shenkin, N. Majalap, J. bin Jami, A. bin Sailim, G. Reynolds, D.A. Coomes, C.J. Chandler, D.S. Boyd, A. Burt, Phil Wilkes, M. Disney, Y. Malhi

## Abstract

The factors that limit the maximum height of trees, whether ecophysiological or mechanical, are the subject of longstanding debate. Here we examine the role of mechanical stability in limiting tree height and focus on trees from the tallest tropical forests on Earth, in Sabah, Malaysian Borneo, including the recently discovered tallest tropical tree, a 100.8 m *Shorea faguetiana*. We use terrestrial laser scans, *in situ* strain gauge data and finite-element simulations to map the architecture of tall broadleaf trees and monitor their response to wind loading. We demonstrate that a tree’s risk of breaking due to gravity or self-weight decreases with tree height and is much more strongly affected by tree architecture than by material properties. In contrast, wind damage risk increases with tree height despite the larger diameters of tall trees, resulting in a U-shaped curve of mechanical risk with tree height. The relative rarity of extreme wind speeds in north Borneo may be the reason it is home to the tallest trees in the tropics.

## Introduction

Tall trees are an essential store of carbon and an inspiration to the public and to ecologists alike (Bastin et al., 2018; Lindenmayer and Laurance, 2016; Lutz et al., 2018). Recently, the world’s tallest tropical tree, a 100.8 m *Shorea faguetiana* (named “Menara”), has been discovered in Sabah, Malaysian Borneo (Shenkin et al., 2019). Trees over 80 m tall have long been recognised in temperate regions - notably the conifer *Sequoia sempervirens* in California and the broadleaf *Eucalyptus regnans* in Tasmania, but the discovery of such tall trees in the tropics is recent. Record-sized tropical trees have also recently been discovered Africa (Hemp et al., 2017) and South America (Gorgens et al. in review).

Tree height growth is driven by the intense competition for light in young forest stands (MacFarlane and Kane, 2017) but why emergent trees continue to grow upwards once they have escaped competition, and what determines the mean and maximum height of forest canopies, is less well understood. Why do the world’s tallest broadleaf trees grow to about 100 m in height, and not 50 m or 500 m?

Possible answers to this question are that tree height is limited by hydraulics, carbohydrate supply or by mechanics. The hydraulic limitation theory posits that the negative water potentials in the vascular systems at the tops of trees, driven by gravity and by demand outstripping supply around midday, inhibits the transfer of water from the vascular system into the cytoplasm of plants (Ishii et al., 2014; Koch et al., 2004). This effect, and the increased risk of embolism, is predicted to lead to lower tree heights in drier regions and is supported by the correlation between canopy height and water availability at the global scale (Klein et al., 2015; Moles et al., 2009a). The carbohydrate transport theory posits that it is the ability of leaves to supply carbohydrates to distant tissues against resistance that limits tree to around 100 m (Jensen and Zwieniecki, 2013), and explains the relationship between water availability and tree height as being determined by climate constraints on leaf size. The mechanical limitation theory suggests that tree height is limited by either gravity, on which most of the literature has focussed due to its tractability, or wind damage risk, which is addressed in the current study. Givnish et al., (2014) showed that the cost of producing and sustaining support tissues increases disproportionately with tree height, meaning that the risk of mechanical damage will also increase. The relative importance of mechanical, hydraulic and carbohydrate constraints will vary according to local climate, and the tallest broadleaf trees are likely to be found in regions with little seasonal water stress, such as aseasonal tropical (Klein et al., 2015; Moles et al., 2009b).

In this study we focus on the mechanical limitation theory in the context of the tallest tropical forests on earth, in northern Borneo. Our key questions are:

1. **How does gravitational stability vary with tree height, architecture and material properties?** Terrestrial laser scanning (TLS) data allow us to estimate the weight of the crowns of tall trees and so test the relative importance of variations in crown size and material properties in terms of their effects on gravitational stability.
2. **How does the risk of wind damage change with tree height?** Taller trees are exposed to higher wind speeds, but they also have better resistance to bending due to their higher trunk diameter and modulus of elasticity. We use field measurements of strain under wind to explicitly test whether these effects balance out.
3. **Is Menara, the world’s tallest tropical tree, close to its mechanical limits?** We focus particular attention on Danum Valley in Sabah, home to Menara and many other tall trees, to illustrate and explore the mechanical limits to tree height.

## Results

**Table 1.** Definitions of key terms.

| Term | Definition |
| --- | --- |
| **Gravitational max height** | The tallest a tree can grow without collapsing under gravity |
| **Modulus of elasticity** | A measure of a material's resistance to bending |
| **K** | The ratio of crown weight to trunk weight |
| **Bending strain** | Extension per unit length – in this case along a tree trunk |
| **Wind-strain gradient** | The slope of the best fit line relating bending strain to squared wind speed |
| **Risk factor** | The measured height of the tree divided by the maximum height consistent with safety, either from wind or gravity |

### Gravitational risk factors decrease with tree height within a forest canopy

We used two models to calculate the gravitational risk factors, the ratio of measured height to theoretical maximum height each tree could reach before buckling under its own weight. The Euler model (equation 1) represents a tree as a tapering cone with realistic trunk diameter and material properties (Greenhill, 1881). The ‘top-weight’ model corresponds to a tapering cone with a top-weight to represent the crown (King and Loucks, 1978). A novel aspect of this study is the inclusion of detailed 3D tree architecture based on TLS data. This allowed us to estimate the woody volume in different parts of the tree and so estimate the weight of the stem and the crown separately.

Both models predicted a decrease in gravitational risk factors with tree height (Figure 2a). This means that, as trees get taller, their radial growth is more than sufficient to compensate for their increased height. We also calculated gravitational risk factors directly from continental height-diameter allometric equations (Feldpausch et al., 2011) and found the same trend (Figure 2b). The intercept and exact shape of these risk factors depend on the relationship between tree height and *K*, the ratio of crown weight to stem weight, but the negative trend and difference between continents do not (S5). This result supports the hypothesis of King et al. (2009) who speculated that understory trees have higher gravitational risk factors due to the prioritisation of vertical height growth under intense competition for light, whereas overstory trees gain less advantage from investing in height growth.

**Figure 1.**
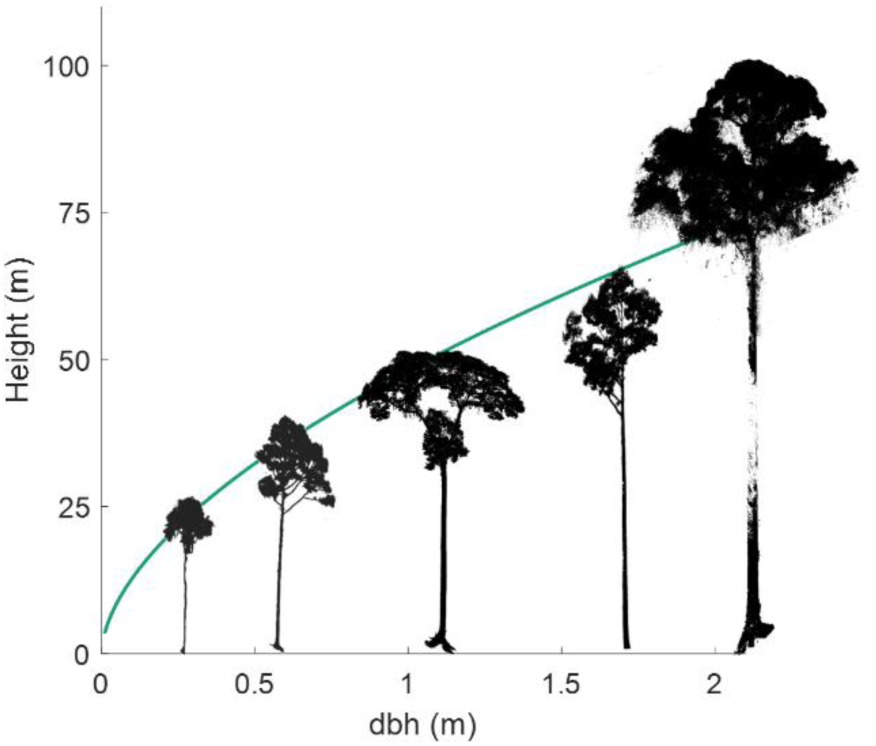
Tree height against trunk diameter measured at breast height (dbh), or above the buttress in buttressed trees, based on the continental allometric equation for Asia (Feldpausch et al., 2011) with trees from this study overlaid. Menara, the world’s tallest tropical tree is at the far right.

**Figure 2.**
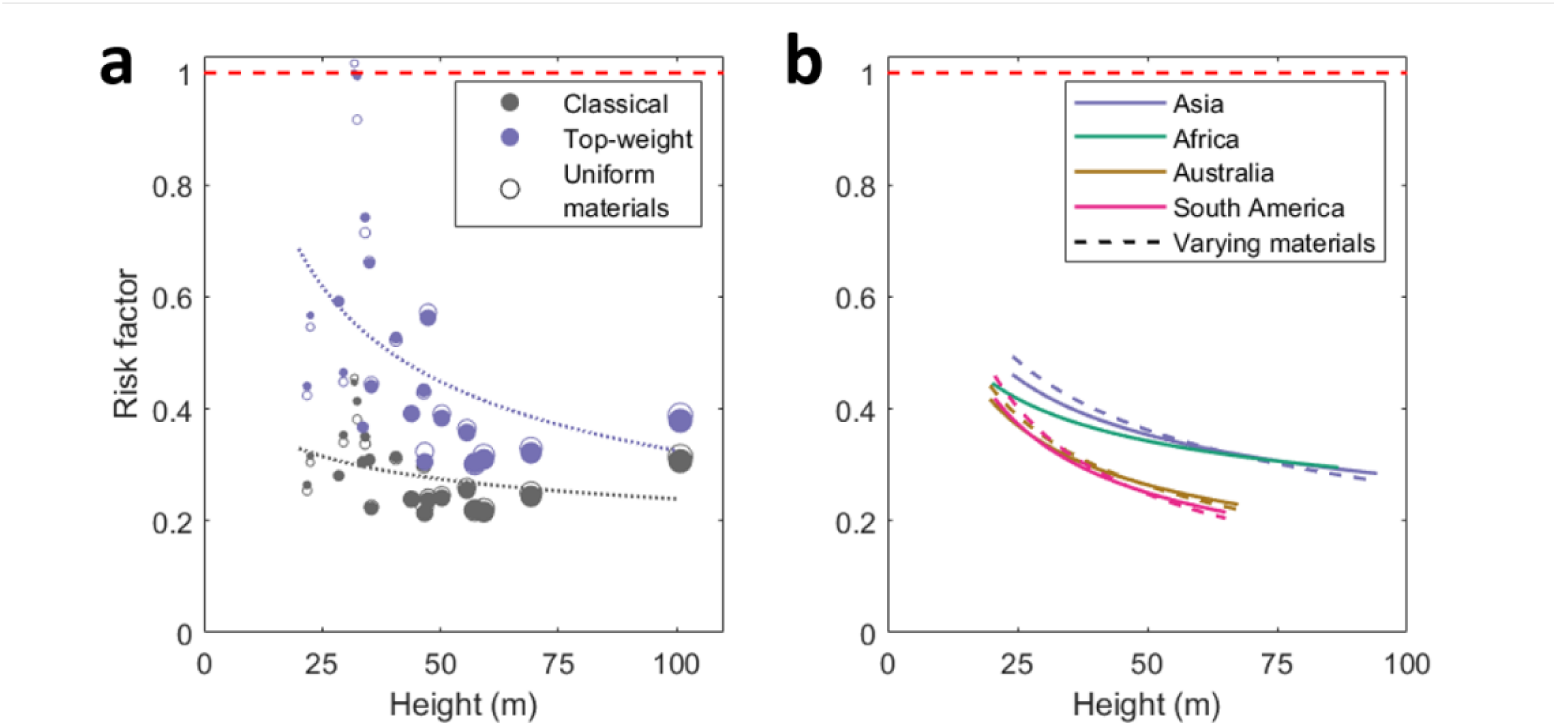
a: Gravitational risk factors for trees in this study calculated using the classical model and top-weight model. The hollow markers represent the same risk factor calculations using the mean material properties of all trees, as opposed to species-specific material properties. Fit lines were calculated based on the latter. b: Gravitational risk factors of trees based on the continental height-diameter allometries using the top-weight model. Solid lines represent uniform material properties while dashed lines indicate material properties varying with height as reported by (Jagels et al., 2018). The red dashed lines at risk factor = 1 shows the point at which a tree would be expected to buckle under its own weight.

### Crown size, not material properties, determine gravitational stability

The risk factor estimates based on the two models followed a similar pattern (adjusted R^2^ = 0.83) but the magnitudes diverged substantially between models (Figure 2a). The ‘top-weight’ model predicted that the trees are much closer to their gravitational stability limit than the ‘classical’ model (increase in risk factor ranged from 0.062 to 0.581 with a mean of 0.214). This means that the presence of a crown substantially decreases overall stability and demonstrates the importance of crown dimensions in gravitational stability.

Comparing continents (Figure 2b) we find that trees in Africa and Asia (the latter data set being dominated by sites in Borneo) have higher gravitational risk factors than those in South America. These differences are driven by tree allometry, not by material properties. Clearly, trees experience similar gravitational forces everywhere on Earth, and we suggest that these differences in gravitational risk factors may be due to differences in wind regime.

Compared to changes in crown size, material properties (i.e. wood density and elasticity) had a substantially lower effect on gravitational stability. Differences in risk factors using species-specific or mean material properties ranged from −0.018 to 0.078 with a mean of 0.004. This small effect of materials on gravitational stability is due to the fact that wood density and elasticity are strongly correlated (Niklas and Spatz, 2010) and it is their ratio which affects gravitational stability.

Although its effect on gravitational stability is small, there is a consistent increase in the ratio of wood elasticity to density with maximum tree height (Jagels et al., 2018). We found that including species-specific material properties (Figure 2a) tended to amplify the difference between short and tall trees, with gravitational risk factors increasing for shorter trees and decreasing for taller trees. Similarly, Figure 2b used the systematic variation in material properties reported by Jagels et al. (2018) and showed a comparable effect.

### Wind risk increases with tree height

The second aspect of mechanical stability is resistance to wind-induced snapping or uprooting (Niklas and Spatz, 2012). Mode-of-death surveys show a wide variation the relative likelihood of snapping and uprooting (Everham and Brokaw, 1996), presumably driven by site conditions, and a survey in Danum Valley found that slightly more trees snapped than uprooted in this site (Gale and Hall, 2001). We do not know of any field technique that can measure the risk of uprooting for large trees in a tropical forest environment. We argue that on average the risk of snapping and uprooting should be similar, as trees are unlikely to have evolved strong mitigation of one risk at the expense of the other (e.g. excessive protection against uprooting would make little sense if tree snapping is the dominant mechanical risk, and *vice versa*). Therefore, we quantified the (more empirically and analytically tractable) risk of snapping and assumed that uprooting is, on average, equally likely.

We measured the local wind speed using anemometers attached to tall trees and measured the bending strains at the base of the trees using strain gauges. As expected, Danum Valley is not a particularly windy site and the maximum recorded wind speed in almost 2000 hours of data collected between August 2016 to March 2017 was a 10 s mean of 7.2 ms^-1^. We selected the maximum wind speed and bending strain for each 10-minute window and calculated the wind-strain gradient (Figures 3a, b). We then calculated each tree’s risk factor for a given maximum wind speed by comparing strain produced at that wind speed with the species-specific breaking strain (Table S1). The absolute value of the wind risk factor depends on the chosen wind measurement type and the aggregation window length. We therefore focus on relative risk factors.

**Figure 3.**
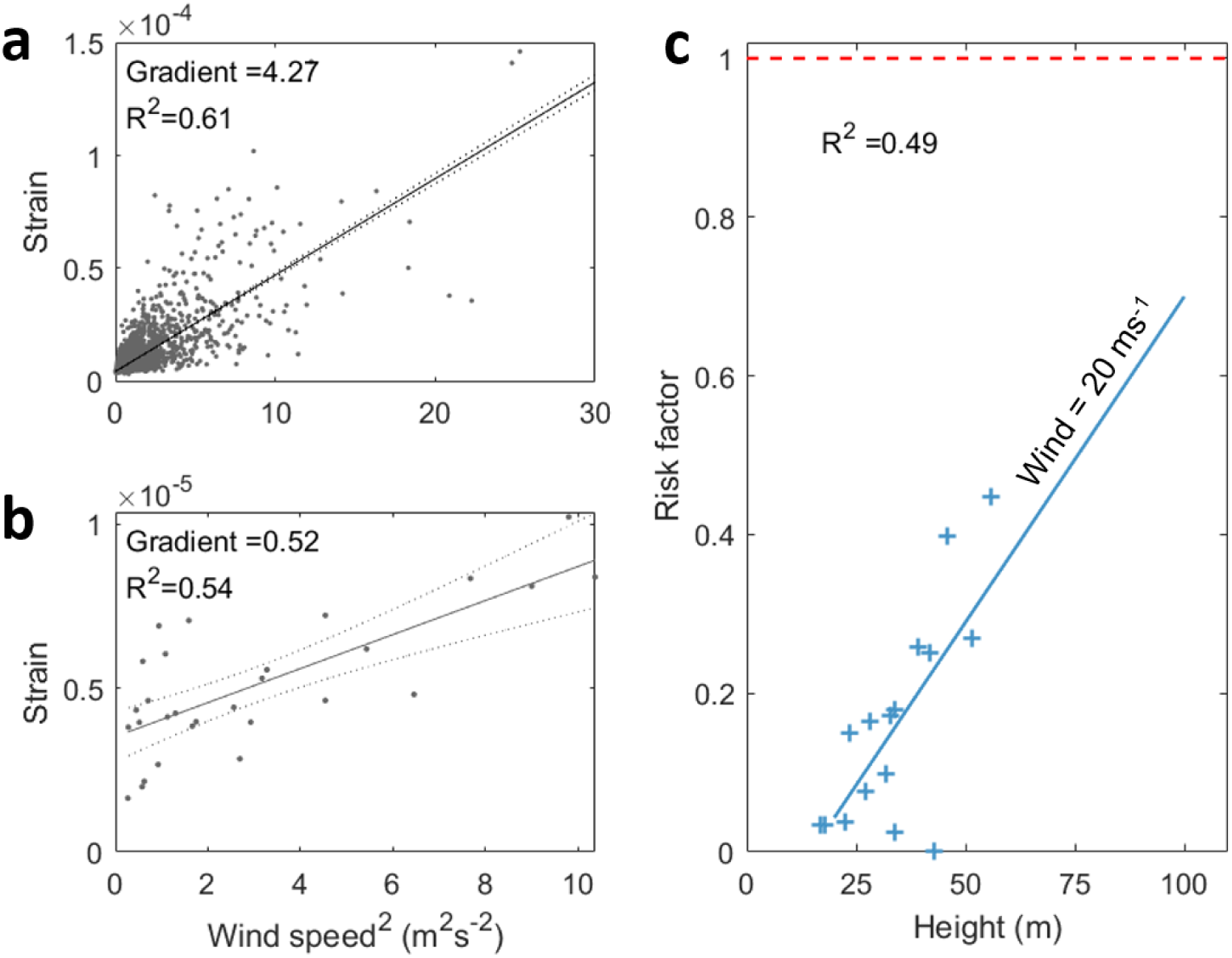
10-minute maximum bending strain against maximum squared wind speed measured with the sonic anemometer for (a) 51 m tall Parashorea malaanonan logged with a CR1000 and (b) 17 m tall Orophea myriantha logged with a CR23X. Note the difference in axes scales and data availability. c: Risk factor against tree height assuming a 20 ms^-1^ wind speed at the position of our sonic anemometer. Individual wind-strain gradients were calculated using a random effects model and a 10-minute aggregation period and risk factors calculated as the ratio of the predicted strain at 20 ms^-1^ wind speed to the breaking strain for that species. For details of the fitting process and sensitivity see S6 and S7. The red dashed line at risk factor = 1 shows the point at which a tree would be expected to break.

The wind-strain gradient was steeper for taller trees (Figure 3c) meaning that taller trees have a higher risk of wind damage. This seems intuitive, since taller trees are exposed to higher wind speeds. However, trees are known to adapt to their local wind environment through increased radial growth (Bonnesoeur et al., 2016; Telewski, 1995). Therefore, it seemed possible *a priori* that these adaptations could counterbalance the increased wind loading. However, our field data demonstrate that this is not the case - the larger diameters of tall trees are not sufficient to compensate for the increased wind loading. This is likely because the support costs scale with trunk circumference and so increase disproportionately with tree height, making it increasingly difficult for a tree to grow radially sufficiently to support the increased wind loading (Givnish et al., 2014). In addition, these results show that variations in material property, such as an increased stiffness for a given density, do not compensate for the increased wind loading.

Post-damage surveys have shown that tall trees are more likely to be damaged by wind than shorter trees (Rifai et al., 2016; Silvério et al., 2018). However, these data represent the first direct measurements of bending strains and mechanistic study of wind damage risk in a tall tropical forest.

### Mechanical risk factor follows a U-shaped curve

We found no significant correlation between the wind and gravitational risk factors for the 11 trees where field and TLS data overlap (p=0.17, 0.08 for the ‘classical’ and ‘top-weight’ models respectively). This demonstrates that overall mechanical stability cannot be approximated by gravitational stability alone. It should be noted that the wind and gravitational risk factors are not simply additive, since the gravitational risk factors are derived from a model whereas the wind risk factors are extrapolated from field data which necessarily include gravitational effects. However, for closed-canopy forests we show clearly that gravity risk factors play the dominant role in the mechanical stability of small trees, while wind damage risk factors dominate for tall trees. The result is a U-shaped mechanical risk profile (Figure 4) illustrating the shift from gravitational limitation to wind limitation as the tree reaches the canopy. Figure 4 also demonstrates that the tree height at which this shift takes place depends on the local wind conditions, with wind damage risk dominating earlier in regions which experience more extreme wind storm events. One hypothesis generated from this work is that the more continental climate system in South America develops larger-scale and stronger storms and squall lines, resulting in shorter average canopy tree height and hence lower biomass storage (Espírito-Santo et al., 2014). Note that it is rare extreme events (e.g. convective downbursts) that would matter, and these rare maximum wind speeds are not necessarily correlated to mean wind speeds.

**Figure 4.**
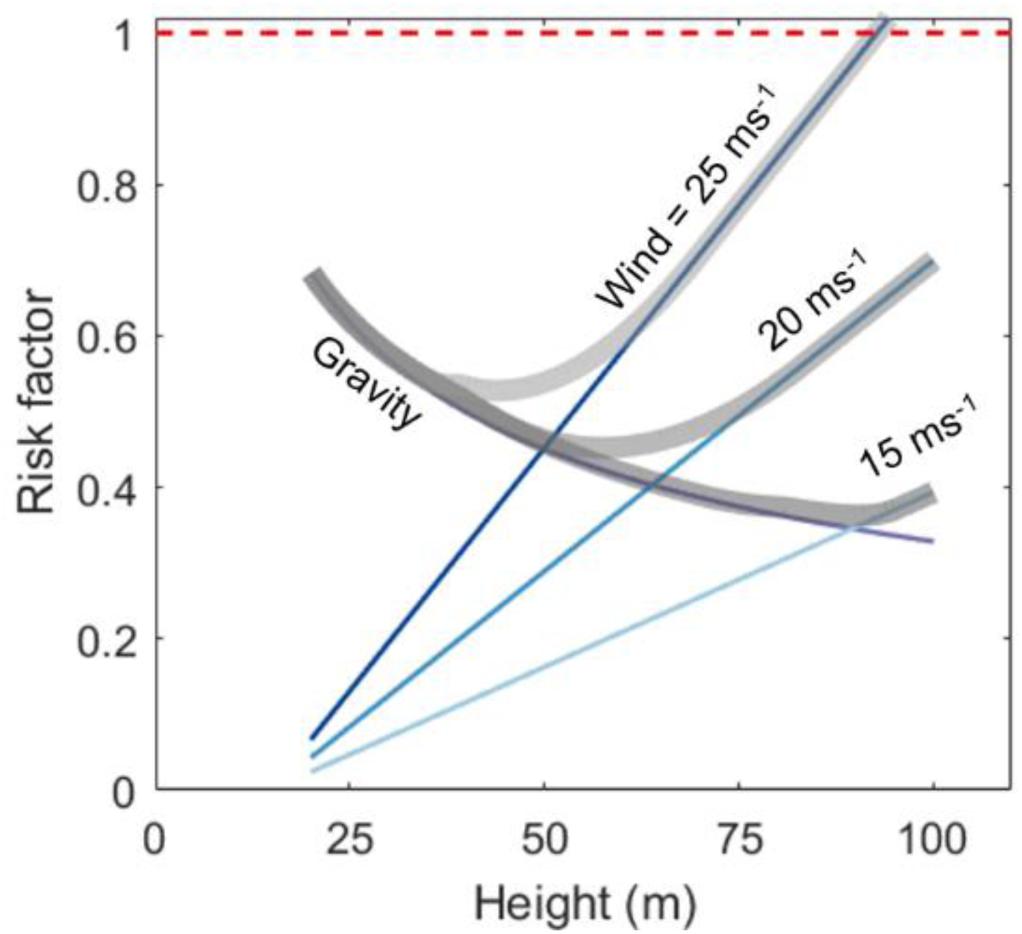
U-shaped variation in mechanical risk factor with tree height. This is derived from the combined gravity and wind risk factors. The red dashed line at risk factor = 1 shows the point at which a tree would be expected to break.

### Is Menara, the world’s tallest tropical tree, close to its mechanical limits?

The world’s tallest tropical tree provides an important case study for questions about mechanical stability. Our analysis shows that the greatest mechanical limitation on this tree’s height is likely due to wind loading. Extrapolating from our field data to the height of this tree we estimate that it would break at 24.1 ms^-1^ (measured at the location of the anemometer used in this study). However, this tree, like the previous record holders from Malaysia (Jucker et al., 2017), is situated near the base of a steep valley and will therefore be sheltered from the wind to a some degree.

In the above analysis we consider gravity and wind separately, but in nature they are combined. In order to simulate the combined effects of wind and gravity we used finite element analysis (FEA), based on a TLS-derived beam model of Menara (Figure 5). FEA is a well-established structural engineering technique and has been successfully applied to conifers (Moore and Maguire, 2008) and temperate broadleaf trees (Jackson et al., 2019; Sellier et al., 2006). Our results demonstrate that the effect of wind dominates the bending response of the tree and that the effect of gravity is secondary (Figure 5). The contribution of gravity increases with wind speed, since the tree deflection is higher and so the overhanging weight of the crown causes a greater moment. Further FEA simulations were carried out for the other TLS-scanned trees, but complications involving modelling gravity for asymmetric trees without information on the asymmetries in material properties make the results difficult to interpret (see S8). However, FEA is a promising technique in tree biomechanics and could be used in future studies to estimate the strength of buttresses, which are characteristic of these giant trees.

**Figure 5.**
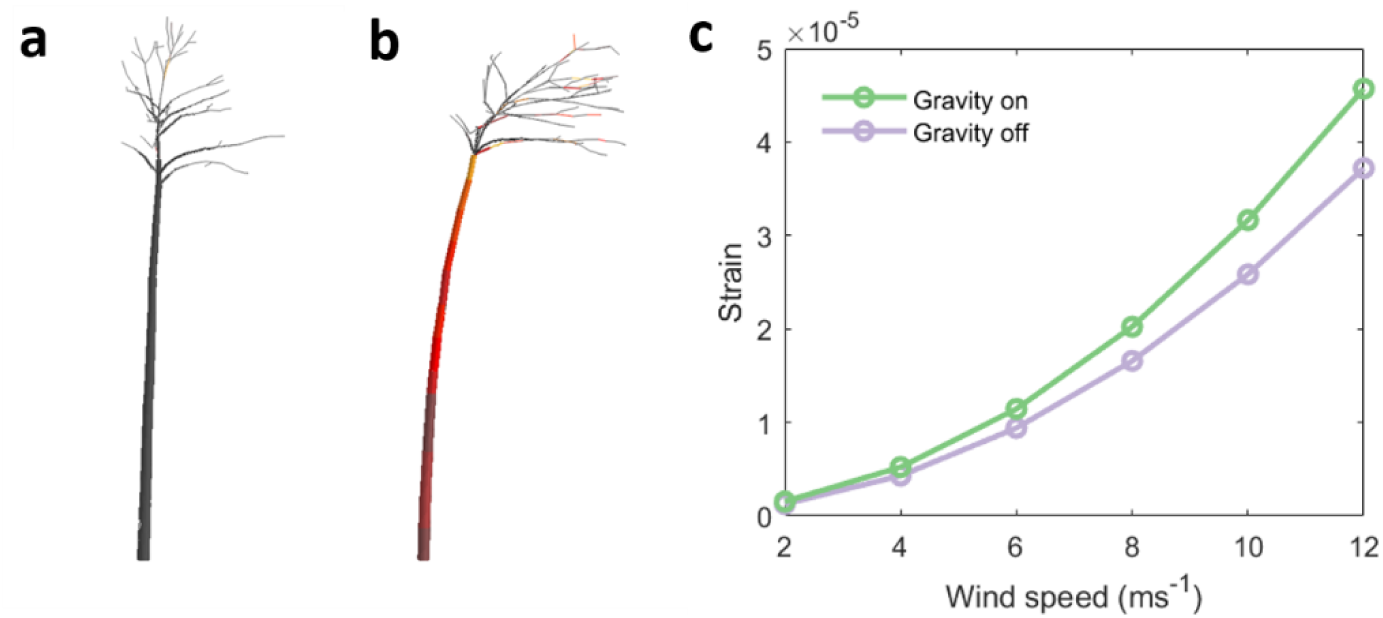
a: Quantitative Structure Model (QSM) of Menara in its rest position. b: Finite element simulation of the same tree under gravity and 12 ms^-1^ wind loading. c: Bending strain produced at the base of the stem of the same tree under simulated wind loading with and without gravity.

## Discussion

We studied the mechanical stability of tall trees in the lowland forests of Sabah, Malaysian Borneo, in order to assess whether these tall trees are subject to mechanical constraints on maximum height. We used TLS data to map the 3D architecture of tall trees and estimated the maximum possible height consistent with gravitational stability. Gravitational risk factors decreased with tree height, meaning that diameter growth more than compensated for increases in tree height, which is consistent with previous work (King et al., 2009; Niklas, 1995). We also found that accounting for the mass of the tree crown substantially reduced the maximum predicted height while material properties had little effect. Our field measurements show that wind damage risk increases with tree height, meaning that neither large diameters nor material properties counterbalanced this risk. Therefore, overall mechanical risk factors follow a U-shaped curve with tree height (Figure 4). The shape of this curve and the point at which mechanical risk shifts from gravity-dominated to wind-dominated depends on the local wind regime.

Overall, this result suggests that wind plays a role in limiting tree height, which has wide-ranging implications including for forest structure and carbon stocks. Two features of geographical variation of tropical tree height support a role for wind constraints. One is the observation that tree heights in Amazonia and Africa are generally lower than in Borneo (Feldpausch et al., 2011). Northwest Amazonia, in particular, has a wet and aseasonal rainfall regime similar to north Borneo (and hence little expected seasonal soil water stress (Malhi and Wright, 2004)), yet much shorter maximum tree heights. We hypothesise that, compared to the insular climate of Borneo, the continental climates of Amazonia or Central Africa are likely to generate more intense convective events and downbursts that lead to a higher frequency of extreme winds. Secondly, at a local scale, we note that the many of the tallest trees in the Danum Valley area, including Menara, appear to be found in somewhat wind-sheltered regions on the lee side of ridges (Shenkin et al., 2019), again suggesting that maximum winds speeds are a limiting factor. The general decline of tree height with increasing dry season intensity does suggest that hydraulic or carbohydrate supply is a constraint on maximum height in many tropical forests, but in forests with little seasonal drought our analysis suggest that rare maximum wind speeds may provide the ultimate constraint.

## Methods

### Description of field sites

The island of Borneo is host to the world’s tallest tropical forests. Danum Valley is situated in the state of Sabah, Malaysia (4°57’N 117°48’E) and is home to a permanent 50 ha forest monitoring plot. Within this plot all the trees with dbh ≥ 1 cm have been tagged and measured and identified to species level where possible. Our study site is located within a 1 ha intense monitoring plot containing 450 trees ≥ 10 cm dbh with a basal area of 32 m^2^, 178 known species and 50 trees whose species are not yet determined (Riutta et al., 2018). Tree height and crown dimensions have also been estimated using a laser rangefinder (Sullivan et al., 2018). The Belian plot in the Maliau Basin Conservation Area (4°44’N 116°58’E), also in Sabah, is a permanent sample plot with trees ≥ 10 cm dbh monitored. It has a basal area of 41.5 m^2^, 135 known species, 4 species determined to genus, and 5 individuals of undetermined taxonomy.

### Terrestrial laser scanning and tree architecture

TLS data were collected in Danum Valley during April 2017 and in Maliau in July 2018 using standard protocols (Wilkes et al., 2017). In this study we focussed on 16 trees from Danum Valley and 5 trees from Maliau, in addition to Menara (Table S1). We manually extracted trees from the plot level point cloud and trimmed the individual tree point clouds manually (Figure S3) using specifically designed software (Vicari, 2017). We used a specifically designed cylinder fitting algorithm (*TreeQSM*; Åkerblom, 2017) to fit 360 Quantitative Structure Models (QSMs) to each tree level point cloud, varying the fitting parameters. The most plausible resulting QSM was selected manually by comparing it to the point cloud (see S2). This process is subjective and all of the point clouds and QSMs are available online. TLS scanning and data processing for Menara followed a similar protocol, except that drone imagery of the crown was used to supplement the TLS data coverage using structure-from-motion techniques (Shenkin et al., 2019). For this tree the QSM of the stem was manually defined using a vertical profile of trunk diameter measurements taken directly from the point cloud. Buttresses were removed from the tree-level point clouds prior to cylinder fitting to generate the QSMs and a cylinder of the same height as the buttress attached to the bottom of the QSM after fitting. The resolution of the point clouds decreased with tree height due to occlusion and the divergence of the TLS laser beam. Therefore, the cylinder fitting process cannot detect the small branches at the tops of the trees and the QSMs systematically underestimated tree height (Figure S2). We therefore use height measurements taken from the point clouds for all analyses except for Menara, which was directly measured by climbing.

As noted by Osunkoya et al., (2007) no generalizable definition of branching order is available in the literature. We therefore manually defined the point at which the crown starts for each QSM (Figure S3). We then calculated the ratio of crown mass to stem mass, *K,* and the positions of the centres of mass for the crown and for the whole tree (Figure S3). Previous work in similar forests found that variation in crown architecture was primarily intraspecific, with only a small proportion of the total variance explained by species (Iida et al., 2011; Osunkoya et al., 2007). These studies also reported a decrease in relative crown size with tree height, which was fit with a power law function (Osunkoya et al., 2007). Therefore, in order to generalize the risk factor calculation, we assumed a power law relationship *K* = 6.92*H*^−0.69^, where the parameters were derived by fitting to the data using the Matlab curve fitting toolbox (Mathworks, 2017). This same relationship was used for all trees in figure 2b so that comparisons between continents or across material properties gradients are not affected by it. The intercept and exact shape of the risk factors depend on the details of this relationship, but the negative trend, difference between continents and effect of material properties do not (S5). It would be interesting to test for systematic variability in this parameter in future work.

### Gravitational stability

The most relevant material properties for mechanical stability are the green wood density, *ρ*_*gw*_, modulus of elasticity, *E*_*gw*_ and modulus of rupture, *MOR*_*gw*_. We collated material properties data from the literature, see S1 for details.

We then calculated the theoretical maximum height each tree could reach before collapsing under gravity, *H*_*max*_, using two different models. The ‘classical’ model (equation 1) of a tapering beam with a circular cross-section (Greenhill, 1881) and the ‘top-weight’ model which includes a top-weight to represent the crown centred at 0.9*H* (King and Loucks, 1978). The difference between the two models is the value of the constant, C. In the top-weight model the weight of the crown is included as per equation 2 which depends on the parameter K, the ratio between stem and crown mass. In the classical model we chose C=1.7464, so that the two models converge when the top weight is set to 0.

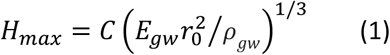

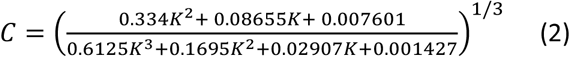

### Wind-strain data collection and processing

Since there is no tower near this plot, anemometers were attached to tall trees (trees 1 and 6) boomed out from the stem. This means that our wind data are site-specific and cannot be translated to the standard 10 m above canopy height point of measurement. However, we found that this unusual method proved highly useful (see S6). In order to measure the tree’s bending in response to wind we attached pairs of strain gauges to the stems of 19 trees at approximately 1.3 m. We used three Campbell Scientific CR1000 data loggers and two CR23X data loggers to record the bending strain data from October 2016 until April 2017. For the seven trees logged with CR1000 data loggers, we collected hundreds of hours of data and found a clear bending strain signal (Figure 2a). We also found clear signals for 9 of the 12 trees monitored with CR23X data loggers, although the data availability was lower and the data were noisier. This complicated the uncertainty analysis and the error propagation for the combined wind-strain gradient against tree height model.

The raw data consist of two mV readings per tree at 4 Hz and we calculated a single maximum strain signal for each tree. This process involved (1) multiplying the raw strain signal by calibration coefficients to transform from units of mV to strain; (2) bandpass filtering the data to smooth out drift (3) re-projecting the signals onto into the North and East facing directions; and (4) combining each pair of strain signals into a single maximum strain signal. We calculated the modal maximum bending strain (Wellpott, 2008) and absolute maximum wind speed for each aggregation period (10 minutes or 1 minute). Using the maxima negates the need for a gust factor (Gardiner et al., 2000) and focuses on the bending strains most relevant to wind damage. We then regressed maximum wind against maximum strain and calculated the gradient. The wind speed measurement system proved useful and wind-strain gradients did not vary much whichever anemometer they were derived from (see S6 and S7).

### Risk factor calculations

We calculated risk factors in order to compare the roles of wind and gravity in maximum height limitation. The gravitational risk factors were defined as the ratio of the measured height (based on the point clouds since the QSMs systematically under-estimate tree height) to the modelled maximum height, *R*_*grav*_= *H/H*_*max*_. We generalized gravitational risk factors using the continental height-diameter allometries (Feldpausch et al., 2011) and, in the case of the top-weight model, the power law relationship between *K* and *H.* We also used variation in material properties for tall and short trees reported by Jagels et al., (2018) to estimate its effect on gravitational stability.

In order to calculate wind damage risk factors, we extrapolated from field data to estimate what the bending strain would be at a chosen threshold wind speed e.g. 20 ms^-1^, *ε*_20_. The risk factor is then *R*_*wind*_= *ε*_20_/*ε*_*break*_, where *ε*_*break*_ is the breaking strain (S1). These risk factors are necessarily relative to the chosen wind speed as well as the point of measurement and aggregating window length. We therefore focus on relative risk factors instead of absolute risk factors. In order to directly compare these results with other sites we would need data on the return time of extreme wind events and wind measurements from towers approximately 10 m above the canopy.

### Finite element analysis of gravity and wind

Finite element analysis is a numerical method used to calculate the stresses and strains over a complex geometry by splitting the problem up into a large number of finite elements. It has been used to simulate the response of trees to wind loading in conifer forests (Moore and Maguire, 2008) and temperate broadleaf forests (Jackson et al., 2019; Sellier et al., 2006). We used the QSM of Menara as the input, and simulated the effect of increasing wind speed both with and without gravity (for details see S8). Simulating the effect of gravity proved difficult, since the trees were obviously scanned in the presence of gravity and are therefore pre-stressed structures. The best approximation to a proper treatment of gravity was to apply a reversed gravity force, export the deformed positions of all the branches into a new analysis, then apply a downwards gravity force and maintain it throughout the simulation. This gave us the desired effect of increasing the moment due to self-weight as the crown deflects under high wind speeds. However, in the cases of other, less straight trees, they failed to stabilize under the downwards gravity load and collapsed. This is likely due to the fact that, in nature, trees develop asymmetric material properties to compensate for their asymmetric architecture, but this variation in material properties cannot yet be included in the simulation.

## Acknowledgments

We thank the Danum Valley Management Committee and Sabah Biodiversity Council for their assistance with fieldwork. TJ was supported by NERC studentship NE/L0021612/1. TJ and DC are supported by NERC grant NE/S010750/1. AS and YM are supported by NERC grant NE/P012337/1 (to YM) and YM is also supported by the Frank Jackson Foundation. DB and CC are supported by NE/P004806/1 and NE/I528477/1. MD was supported in part by NERC NCEO for travel and capital funding for lidar equipment, and NERC Standard Grants NE/N00373X/1 and NE/ P011780/1. Some TLS data collection was funded through the Metrology for Earth Observation and Climate project (MetEOC-2), grant number ENV55 within the European Metrology Research Program (EMRP). The EMRP is jointly funded by the EMRP participating countries within EURAMET and the European Union. Some TLS data was contributed by the NASA Carbon Monitoring System, NASA Future Mission Fusion for Improving Aboveground Biomass Estimation in High Biomass Forests

## Data availability

The field data collected for this study are available at (https://doi.org/10.5285/657f420e-f956-4c33-b7d6-98c7a18aa07a). The field data processing scripts, summary data, TLS point clouds and QSMs are all available (https://github.com/TobyDJackson/WindAndTrees_Danum). Radius and mass taper exponents were calculated in Matlab (https://github.com/TobyDJackson/TreeQSM_Architecture). An updated library for analysing tree structural information is also available in R (https://github.com/ashenkin/treestruct).

