## Supplementary material for "The mechanical stability of the world’s tallest broadleaf trees"

1 – Environmental Change Institute, School of Geography and the Environment, University of Oxford, OX1 3QY, UK
2 – Forest ecology and conservation group, Department of Plant Sciences, University of Cambridge, CB2 3EA, UK
3 – Phytochemistry Unit, Forest Research Centre, Jalan Sepilok, 90715 Sandakan, Sabah, Malaysia
4 – South East Asia Rainforest Research Partnership (SEARRP), Danum Valley Field Centre, 91112 Lahad Datu, Sabah, Malaysia
5 – School of Geography, University of Nottingham, Nottingham, NG7 2RD, UK
6 – Department of Geography, University College London, WC1E 6BT, UK
7 – NERC National Centre for Earth Observation (NCEO), Leicester, UK

### *S1. Material properties overview table*

| # | Site | Tree tag | TLS ID | Genus & species | DBH cm | Height m | $\rho_{gw}$ kgm$^{-3}$ | $E_{gw}$ MNm$^{-2}$ | $MOR_{gw}$ MNm$^{-2}$ |
|---|---|---|---|---|---|---|---|---|---|
| **1** | DV | 30343 | A1 | *Parashorea malaanonan* | 110 | 51 | 641 | 9100 | 61 |
| **2** | DV | 30296 | B2 | *Chisocheton -* | 29.8 | 23 | 626 | 7443 | 53 |
| **3** | DV | 30281 | C3 | *Pometia pinnata* | 32.2 | 28 | 787 | 9399 | 69 |
| **4** | DV | 30313 | D4 | *Shorea parvifolia* | 55 | 39 | 513 | 6700 | 48 |
| **5** | DV | 30476 | E5 | *Aporusa benthamiana* | 23.3 | 24 | 781 | 9326 | 68 |
| **6** | DV | 10498 | F6 | *Shorea parvifolia* | 73 | 46 | 513 | 6700 | 48 |
| **7** | DV | 10542 | G7 | *Terminalia citrina* | 49.2 | 42 | 961 | 13000 | 94 |
| **8** | DV | 20447 | H8 | *Parashorea malaanonan* | 100 | 56 | 641 | 9100 | 61 |
| **9** | DV | 20445 | | *Orophea myriantha* | 20.6 | 17 | 747 | 9680 | 70 |
| **10** | DV | 20419 | I10 | | 39.4 | 31 | 747 | 9680 | 70 |
| **11** | DV | 30409 | J11 | *Caryodaphnopsis tonkinensis* | 43.5 | 39 | 747 | 9680 | 70 |
| **12** | DV | 10676 | K12 | *Shorea pauciflora* | 24.1 | 34 | 689 | 9700 | 68 |
| **13** | DV | 20523 | L13 | *Syzygium panzer* | 40.2 | 33 | 798 | 9531 | 70 |
| **14** | DV | 10612 | | *Diospyros tuberculata* | 35.3 | 34 | 567 | 6728 | 47 |
| **15** | DV | 20557 | M15 | *Canarium pilosum* | 29.9 | 32 | 593 | 6200 | 41 |
| **16** | DV | 20596 | | | 21.1 | 27 | 747 | 9680 | 70 |
| **17** | DV | 20593 | N17 | *Eusideroxylon zwageri* | 73.3 | 43 | 1282 | 17700 | 143 |
| **18** | DV | 10643 | | *Orophea myriantha* | 18.6 | 18 | 747 | 9680 | 70 |
| **19** | DV | 20649 | | *Eusideroxylon zwageri* | 35.4 | 22 | 1282 | 17700 | 143 |
| **20** | DV | 50214 | O20 | *Parashorea malaanonan* | 115.0 | 52 | 641 | 9100 | 61 |
| **21** | DV | 20629 | P23 | *Shorea johorensis* | 135.0 | 53 | 689 | 9700 | 68 |
| **22** | DV | Menara | | *Shorea faguetiana* | 2.13 | 100.8 | 673 | 9700 | 66 |
| **23** | MLA | 102 | MLA_157 | *Shorea gibbosa* | 171.2 | 69.2 | 625 | 9000 | 67 |
| **24** | MLA | 60 | MLA_215 | *Shorea faguetiana* | 162.9 | 59.2 | 673 | 9700 | 66 |
| **25** | MLA | 152 | MLA_38 | *Shorea dasyphylla* | 110.2 | 46.7 | 609 | 9700 | 63 |
| **26** | MLA | 158 | MLA_39 | *Payena microphylla* | 92.7 | 43.9 | 700 | 9356 | 67 |
| **27** | MLA | 195 | MLA_40 | *Shorea parvifolia* | 157.8 | 57.3 | 513 | 6700 | 48 |

*Table S1 – Overview of the trees in this study. $\rho_{gw}$ is the green wood density, $E_{gw}$ is the green wood elasticity and $MOR_{gw}$ is the green wood modulus of elasticity.*

Niklas and Spatz, (2010), collated measurements of green wood properties and we preferentially use measurements from this data set. Two trees could not be identified to genus level and were assigned the mean material properties of the other trees. Species-specific material properties data were available for 6 of the 19 focal trees. Where no species-specific material properties information was available, we used a genus level average. This covered a further 6 trees, but for 5 identified trees there was no species or genus level information available in (Niklas and Spatz, 2010). In these cases, we used the Global Wood Density data base (Chave et al., 2009) to look up the dry wood density, $\rho_{dw}$, taking an average where there were multiple samples of the same species or a genus average where no species-specific information was available. We then used the data set compiled by Lavers (1983) to relate dry wood density to green wood density and elasticity (equations 3-5). The Lavers (1983) data set is a subset of the Niklas and Spatz (2010) data set, except that it also gives dry wood density for each species as well as uncertainty estimates. Full details of the material properties used are given in Table S1. We found the following relationships for N=178 hardwoods:

$$\rho_{gw} = 1.2 \times \rho_{dw} + 27.3 \ \ (R^2 = 0.97, RMSE = 36) \quad (3)$$

$$E_{gw} = 14.6 \times \rho_{dw} + 157.6 \ \ (R^2 = 0.72, RMSE = 1588) \quad (4)$$

$$MOR_{gw} = 0.12 \times \rho_{dw} - 7.14 \quad (\mathrm{R}^2 = 0.78, \mathrm{RMSE} = 11) \quad (5)$$

### S2. Terrestrial laser scanning data processing

We found a good agreement between manual and QSM measurements of dbh, but a systematic underestimation of tree height from the QSMs relative to the point clouds (Figure S2), due to the beam divergence causing increasing noise at the tops of the trees. The overall size of the crown estimated from the QSMs followed a similar pattern to those estimated from the ground. There were large differences, which is unsurprising given that these measurements were not defined in the same way, but these errors were normally distributed.

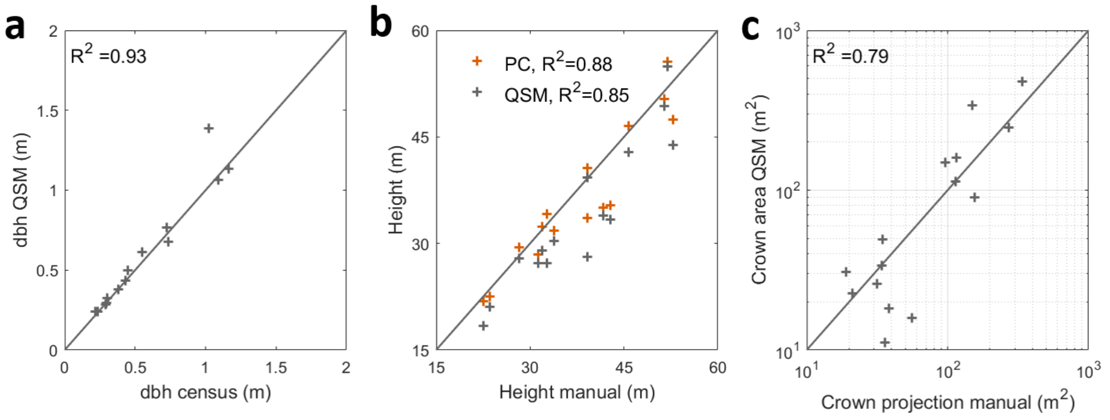


*Figure S2 – Field data against QSM measurements for (a) dbh (b) height and (c) crown dimensions. Note that the two crown dimension variables are not identical and this plot is presented on a log-log scale.*

### S3. Describing tree architecture using terrestrial laser scanning data

We measured tree diameter at multiple heights directly from the point clouds and used these data to calculate the radius taper exponent, $\alpha$

$$r(z) = r_0\left(\frac{H-z}{H}\right)^{\alpha} \quad (6)$$

where $r(z)$ is the radius at height z, $r_0$ is the radius at the base of the trunk and $H$ is the tree height. In order to calculate the mass taper exponent, $\beta$, we split the QSM into vertical segments and summed the volume of the cylinders within each segment (Figure 6C). We assume uniform density and fit a relationship of the same form as equation 6. Similar to the previous models, the tree is modelled as tapering beam with the same diameter as the tree, but in this case the taper exponent $\alpha$ is specific to each tree. Instead of a top weight, this crown is modelled as a distributed mass with a functional form similar to the radius taper (equation 6) and a specified total mass.

We also calculated the vertical profile of sail area, $A_{sail}(\mathrm{z})$, for each tree based on the QSM in two ways. First, we calculated the total area of the cylinders projected into the X-Z plane. Second, we calculated the total width of the tree at each height. We calculated these widths from eight points of the compass and took used the mean sail profile (Figure S3).

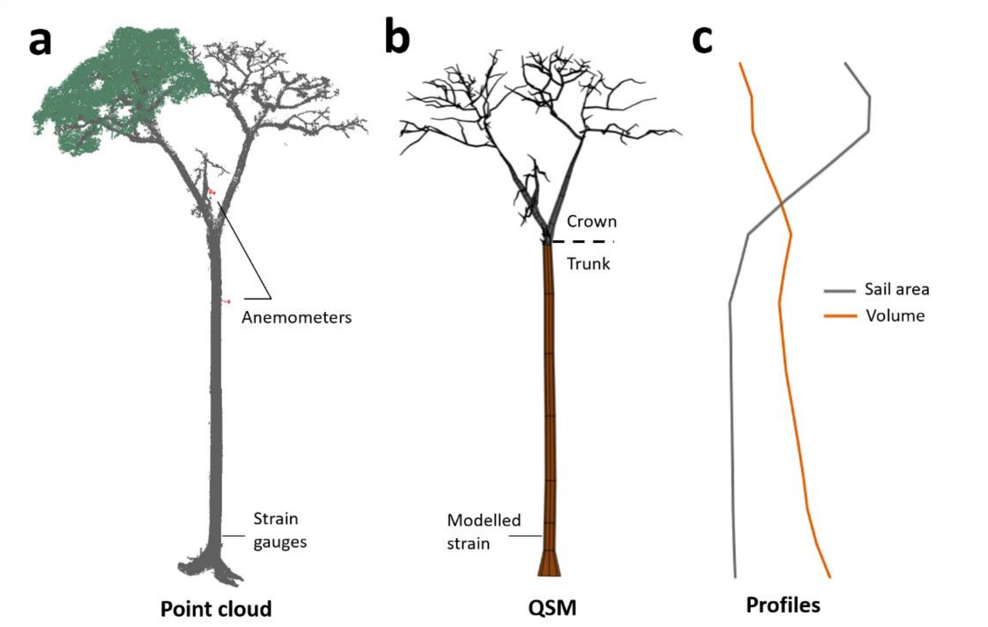


*Figure S3 – A: point cloud of tree 1 (height = 51 m, dbh = 110 cm), showing the position of the anemometers and strain gauges. This image also gives an example of the manual leaf filtering, with points assumed to be leaves shown in green on part of the crown. B: Simplified QSM for tree 1, showing manual definition of the crown and the centre of volume for the whole tree (lower) and the crown (upper). C: Vertical profiles of sail area and volume derived from the QSM.*

### *S4. Tree specific gravity model*

In addition the two models of gravitational stability presented in the main text (Greenhill, 1881; King and Loucks, 1978), we also tested a third model. This model was adapted from Greenhill (1881) by Jaouen et al., (2007) . The maximum height is given by

$$H_{max} = \sqrt{\frac{\pi E_{gw}}{16 M_{tot} g}} \cdot r_0^2 \cdot b \cdot (|\beta - 4\alpha + 2|) \qquad (7)$$

Where $M_{tot}$, is the total mass of the tree and $b$ is the first root of the Bessel function $J_x$ with

$$x = (4\alpha - 1)/\,(\beta - 4\alpha + 2) \qquad (8)$$

Based on this model, four of the shorter trees actually exceed their predicted maximum height (Figure S4). This suggests that many of the shorter trees in this sample could not safely grow much taller with their current mass and radius taper exponents or alternatively that the model underestimates their maximum height. We used this model to compare the gravitational stability of our tall broadleaf trees with Californian *Sequoia sempervirens*, the tallest conifer species on Earth. Comparing the classic model and the tree specific model (Figure S4) shows that, whereas the shape of the broadleaf trees brings them nearer the gravitational limit, the shape of the conifers has a much smaller effect. This is because the crown mass in conifer trees is distributed along the stem and the centre of mass of the crown is much lower, meaning that the tree is more stable. These conifers can grow up to 120 m, whereas the tallest broadleaf trees ever described are approximately 100 m.

We did not use this tree-specific model in the main text because we found it to be sensitive to the mass taper exponent, which was strongly dependent on the QSM quality. Furthermore, the interpretation of risk factors based on this model is more complex due to the fact that the model is specific to each tree. In the case of the classical model, the risk factor compares how close each tree is to the maximum height of a tapering cone of the same diameter. In this case, the risk factor compares how close each tree is to the maximum height of a cone with the same diameter and total mass, whose stem radius taper and mass distribution are the same as the tree. It is unclear whether stem radius taper changes with ontogeny, but it seems likely that both mass distribution and total mass would change. Thus the risk factor quantifies the height that such a tree could have reached in optimal conditions with the same investment of resources. This model of gravitational stability was not included in the main text for these reasons.

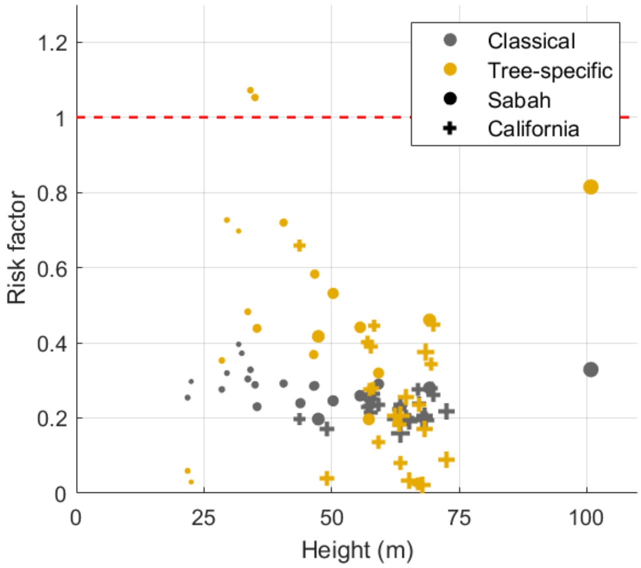


*Figure S4 – Gravitational risk factors for trees from Sabah and California calculated using the classical model and the tree-specific model of* (Jaouen et al., 2007).

### *S5. Sensitivity of the gravitational risk factor calculation to the ratio of crown volume to stem volume*

The gravitational risk factor calculation using the top-weight model requires a parameter, K, defined as the ratio of crown volume to stem volume, assuming a constant wood density. This is a measure of how top-heavy the tree is, which will obviously have a significant effect on its stability. We can estimate K from TLS data for the focal trees in this study. However, in figure 2b we seek to generalise across continents using allometric equations and we did not possess any data on K at this scale. We therefore found the most representative relationship between K and tree height (figure S5c) for our small sample of trees and applied it to all continents (figure S5f and main text figure 2b). This simplification could be improved upon by collating destructive sampling data and TLS data for as many tropical sites as possible and testing whether there are systematic differences in the variation of K with tree height, but this was beyond the scope of this study.

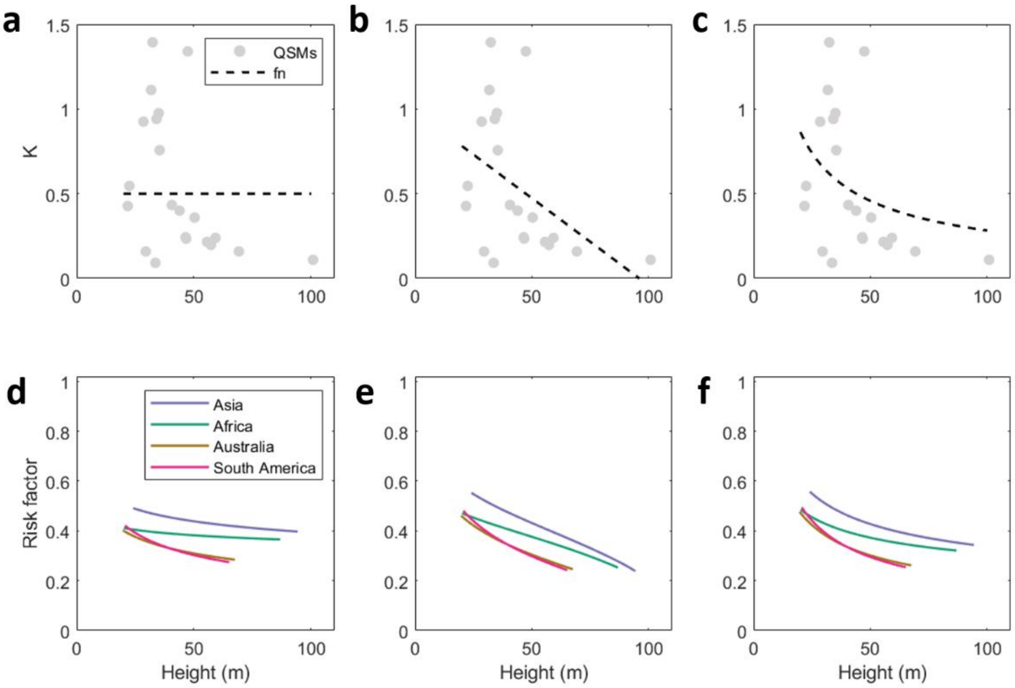


*Figure S5 – Top row: Ratio of crown volume to stem volume, K, against tree height fitted with (a) single value (b) linear function and (c) power law. Bottom row: Gravitational risk factor against tree height for continental allometries. These calculations are all based on the top-weight model and the function for K in the panel directly above i.e. (d) single value (e) linear function and (f) power law.*

### *S6. Measuring wind from tall trees*

One limitation of this study was the lack of an above canopy tower from which to measure wind speed. Instead, we attached anemometers to tall trees and measured the wind speed above the dense canopy but below the canopy of the emergent trees. This means that the wind measurements are site-specific.

Cup anemometers were mounted at 25 m and 35 m on tree 1 (Figure S3). In the case of tree 6 a cup anemometer was mounted at 19 m and a 2D sonic anemometer at 30 m. The cup anemometers were Vector Instruments A100LK/5M and recorded wind speeds at 0.1 Hz, and the 2D sonic anemometer was a Gill Sonic 1, recording at 1 Hz. This method of attaching anemometers to emergent trees is unusual and best practice is to measure measured wind speeds above the canopy. However, this requires a tower which greatly limits the choice of study site. A professional tree climber attached the anemometers to steel poles, which were boomed out from the tree by 1.5 m and supported above and below. The anemometers were attached pointing in different directions to each other so that any directional bias would be evened out. Four anemometers were mounted in this way, at two different heights on two trees which were 63 m apart. This setup was successful insofar as the wind-strain relationships did not vary substantially depending on which anemometer was used. However, this unusual method of measuring wind speed means that our results would not be directly comparable with other sites nor with above canopy wind measurements.

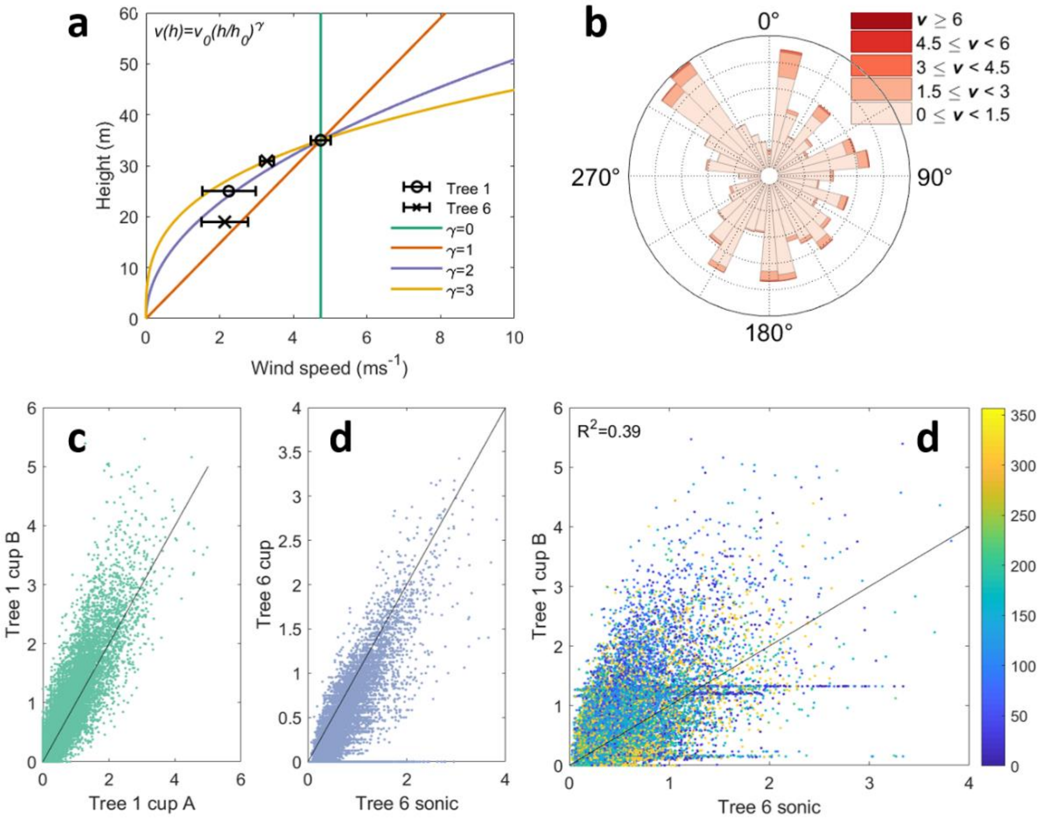


*Figure S6 – A: vertical profile of wind speed measurements with possible power law fits overlaid. B: wind rose for 10-minute maximum wind speeds from the sonic anemometer. C and D: 10-minute maximum wind speeds measured at two heights in the tree 1 and 6, respectively. E: Correlation between 10-minute maximum wind speeds measured at tree 6 and that at tree 1, 63 m away, coloured by wind direction.*

### *S7. Wind-strain gradients*

The data loggers were placed at the base of tall trees, selected for height and accessibility. To minimize the length of wire used for each strain gauge, subsequent trees were selected nearby. The strain gauges were attached to the data loggers via wheat-stone bridge circuits to balance the inevitable offset in voltage. An excitation voltage was pulsed through the circuit at 4 Hz and the voltage across the strain gauges monitored at the same frequency. The limited storage capacity of the CR23X data loggers was supplemented using a Raspberry Pi, as in (Jackson et al., 2019). To reduce the power used by the Raspberry Pi, a trigger system was implemented so that it would only turn on and record data when the variance of the bending strain signal was high (i.e. windy periods). Unfortunately, the voltage offset increased further than expected due to temperature and humidity variations, and this led to a reduced data logger resolution, the failure of the trigger system and ultimately to loss of data. Nevertheless, we recorded some data for these CR23X logged trees and the CR1000 data loggers were unaffected by this, since they had sufficient internal storage.

Equipment limitations led to the loss of some data for 12 of the 19 trees. Nine of these trees retained sufficient usable data and appear in Figure 3, but the reduced data availability led to lower confidence in the derived wind-strain gradients and hampered uncertainty analysis. This meant that, while the broad trend was clear, the field data was error prone. We pooled the data and fitted a random slope model of the form $strain{\sim}v^2 + (v^2 - 1|tree)$. This allows the slope of the wind-strain relationship to vary for each tree but keeps the intercept fixed for all trees.

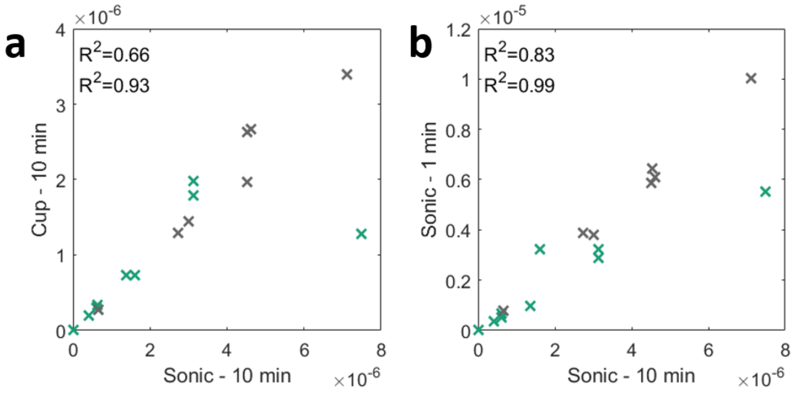


*Figure S7 – Sensitivity of wind-strain gradients to data processing method. All of the axes are wind-strain gradients with units $\mu m^{-2} s^2$. The x-axes are identical both panels - the baseline wind strain gradient as used in the main text, which was calculated using a 10-minute aggregation period and data from the sonic anemometer. The y-axes are wind-strain gradients using (a) wind speed data from a cup anemometer 63 m away from the sonic anemometer (b) a one-minute aggregation period. Note the difference in y-axis ranges. The first $R^2$ values refer to all of the trees in the graph while the second refers to only those trees logged with the CR1000s.*

### *S8. Finite element analysis of tall trees*

We used finite element analysis to simultaneously model the tree bending in the wind and the increasing moment caused by the weight of the overhanging crown. In order to prepare for finite element analysis we simplified the chosen QSM by averaging together neighbouring cylinders and removing all cylinders with a radius under a given threshold. This threshold was changed based on the specific QSM for each tree, since taller trees had a lower point density in their crowns. Each simplified QSM was imported into Abaqus (Simulia Software Company, 2017) via the input file scripting language, using freely available software (Jackson, 2017). Each cylinder was modelled as a Euler-Bernoulli beam (B31) and assigned either the mean material properties or the species-specific properties given in Table S1. Point clouds, original QSMs and simplified QSMs are available online (https://github.com/TobyDJackson/WindAndTrees_Danum)

In the field, a tree's resting position is the equilibrium position between gravitational forces and stresses in the wood. In engineering terms, trees are pre-stressed structures. In order to model the effect of gravity accurately, we therefore first apply a reversed gravitational force to each tree, causing the branches to bend upwards. We then extracted the deformed position of each branch and imported these into a new Abaqus session. A gravitational force was then applied downwards, causing the branches to return to their initial position but now with stresses and strains from the trees self-weight. This gravitational force was maintained throughout the analysis, so that if the tree bends in the wind, the additional force of the overhanging crown is modelled. In some cases where the tree was leaning or the crown highly asymmetric the tree passed its original position and continued to deflect unrealistically.
